# Functional characterization of a small gene family coding for putrescine hydroxycinnamoyltransferases in tomato

**DOI:** 10.1101/2022.12.17.520863

**Authors:** Marwa Roumani, Sébastien Besseau, Alain Hehn, Romain Larbat

## Abstract

Phenolamides are specialized metabolites widely distributed in the plant kingdom. Their structure is composed by the association of hydroxycinnamic acid derivatives to mono-/poly-amine. This association is catalyzed by N-hydroxycinnamoyltransferases enzymes. Tomato plants are accumulating putrescine-derived phenolamides in their vegetative parts. Recently, we identified two genes coding for putrescine-hydroxycinnamoyltransferase (PHT, *Solyc11g071470 and Solyc11g071480)*, which control the accumulation of caffeoylputrescine in tomato leaf submitted to the infestation of leafminer. In this study, we prospected for additional genes implicated in the accumulation of putrescine-derived phenolamides in the tomato vegetative organs. We identified two genes (*Solyc06g074710* and *Solyc11g066640*) that we functionally characterized as new PHT. The substrate specificity and the expression pattern *in planta* was determined for the four tomato PHT. Taken together the results give a comprehensive view of the control of the putrescine-derived phenolamide accumulation in tomato plant through the biochemical specificity and the spatial expression of this small family of PHT.

**Main conclusion:** We identified and functionally characterized two new putrescine hydroxycinnamoyl transferases (PHT) in tomato. These enzymes complete a set a four PHT which control the distribution of putrescine-derived phenolamides in tomato plants.

## Introduction

Phenolamides are specialized metabolites widely distributed in plants. They result from the association of phenolic acids and mono- or poly-amines (tyramine, dopamine, serotonin, putrescine, agmatine, spermidine, spermine) which leads to a high structural diversity (for review, see Bassard et al. 2010; Macoy et al. 2015; Roumani et al. 2021). Phenolamides are reported in flowers, seeds and pollen coat, but their biological role in these organs are not elucidated yet (Cabanne et al. 1981; Martin-Tanguy 1985, 1997; Vogt 2018). In Solanaceae, Brassicaceae and Poaceae, these metabolites are also found in vegetative organs. They are accumulating in response to abiotic stresses (Matsuda et al. 2005; Demkura et al. 2010) and biotic stresses such as pathogen infection (Roepenack-Lahaye et al. 2003a; Zacarés et al. 2007; Muroi et al. 2009; Royer et al. 2016a; Morimoto et al. 2018; Ube et al. 2019) or herbivory (Pearce et al. 1998; Tebayashi et al. 2007; Kaur et al. 2010a; Onkokesung et al. 2012a; Marti et al. 2013; Alamgir et al. 2016). In these latter cases, phenolamide accumulation was proved to contribute to the plant defense against bioagressors through direct antimicrobial activities (Zacarés et al. 2007; Yogendra et al. 2015; Dobritzsch et al. 2016; Yogendra and Kushalappa 2016; Ube et al. 2017; Morimoto et al. 2018), reinforcement of the plant cell wall (King and Calhoun 2005), perturbation of the insect oviposition (Tebayashi et al. 2007) and insect toxicity leading to an increase in mortality or at least impacts on the life cycle (Kaur et al. 2010a; Onkokesung et al. 2012a; Marti et al. 2013; Alamgir et al. 2016).

The biosynthesis of phenolamides involves N-hydroxycinnamoyltransferases that transfer the phenolic acid donors to the mono- or poly-amine acceptors. To date, about fifty N-hydroxycinnamoyltransferases have been biochemically characterized (Wang et al. 2021; Roumani et al. 2021). They belong to two different enzyme classes. Hence, some of them are homologous to GNATs (Gen5-related N-acetyltransferases), but most of them belong to the BAHD acyltransferase superfamily. GNATs are restricted the biosynthesis of monoamine phenolamides (tyramine, dopamine, noradrenaline, octopamine). BAHD N-hydroxycinnamoyltransferases have a much broader activity and are able to catalyze the biosynthesis of a large diversity of phenolamides. Depending on their amine specificity and their phylogeny, we recently proposed to classify this BAHD N-hydroxycinnamoyltransferases into eight clusters (Roumani et al. 2021).

*Solanum lycopersicum* has been reported to produce constitutively several phenolamides. For examples, two- and three-substituted spermidine derivatives were detected in pollen coat (Paupière et al. 2017) and putrescine derivatives are found in healthy vegetative organs (Larbat et al. 2014). This chemodiversity is significantly increased when the plants are subjected to biotic stresses leading to the synthesis of dopamine, tyramine, noradrenaline, octopamine and, more recently, tri-substituted spermine derivatives accumulating locally and/or systemically (Von Roepenack-Lahaye et al. 2003; Zacarés et al. 2007; Lopez-Gresa et al. 2016; Roumani et al. 2022). Interestingly, in comparison with this diversity, only a few N-hydroxycinnamoyltransferases have been reported in this plant, among them, a 4-members family of enzymes using tyramine/noradrenaline as acceptors (Roepenack-Lahaye et al. 2003) and one catalyzing tri-acylation of spermidine (Perrin et al. 2021). Recently, two genes encoding putrescine-hydroxycinnamoyltransferase (PHT) were reported (*Solyc11g071470/ Solyc11g071480)*. The expression of these genes was described to be dramatically induced in response to herbivory and to contribute to an increase of the accumulation of caffeoylputrescine in the damaged tissues (Roumani et al. 2022). Such genes coding for PHTs were already described in *N. attenuata* (Onkokesung et al. 2012), in Arabidopsis (Wang et al. 2021) and in rice where this activity was also supported by a small gene family displaying complementary expression profile (Dong et al. 2015; Tanabe et al. 2016).

The aim of this study was to characterize the determinants of putrescine-derived phenolamide accumulation in tomato plants. We identified two additional genes encoding PHTs. The biochemical properties and the expression profiles of the four enzymes were investigated with regard to the phenolamide accumulation *in planta*.

## Materials and methods

### In silico investigations and phylogenetic analysis

The protein sequences of *Solyc11g071470* and *Solyc11g071480* previously functionally characterized as putrescine hydroxycinnamoyltransferase (Roumani et al. 2022) were blasted against a tomato genome protein sequence database (ITAG 3.20) using the BLASTp procedure. Proteins showing more than 60% identity were taken into account for the rest of the paper.

A phylogenetic tree was constructed with the protein sequences identified in tomato and a selection of BAHD acyltransferases functionally characterized, and representing the 8 clades of N-hydroxycinnamoyltransferases within this enzyme family (Roumani et al. 2020). Protein sequences were aligned with ClustalW and the tree was further constructed using neighbor-joining method with bootstrap values corresponding to 100 resampling.

The 3000 bp upstream sequence from the ATG initiation codon of each gene were used for promoter sequence analysis according to Chen et al. (2021). The genetic distance between the 4 promoters was calculated using MEGA 11 software. In addition, the four promoter sequences were searched for regulatory elements using the New PLACE database (https://www.dna.affrc.go.jp/PLACE/?action=newplace).

### Plant material, growth conditions and treatments

Tomato plants (*Solanum lycopersicum*, var. Better Bush) were grown from seed in hydroponic conditions in a growth chamber (16 h photoperiod, 23°C/18°C day/night, 60 % air humidity). The nutrient solution, with a nitrogen concentration at 7 mM, was prepared with water and pure salts as described in supplementary data 1.

For the spatio-temporal study of the phenolamide accumulation and tomato gene expression, three harvests were realized. A first one on 10-day-old plantlets, which were separated into three organ classes: Rootlets, stem and cotyledons. A second harvest was done on 28-day-old vegetative plants, corresponding to 6 leaves-plants which were separated into 5 organ classes: Roots, “Low Stem” corresponding to a stem section between leaf 1 and leaf 4, “High Stem” corresponding to stem section between leaf 4 and leaf 6, “Mature Leaves” corresponding to the limb part of leaves 3 and 4, and “Young Leaves” corresponding to the limb of leaves 5 and 6. A last harvest on 40-day-old flowering plants was focused on flowers by discriminating “Flower Buds”, “Closed Flowers” and “Open Flowers”.

For the inducible study of the phenolamide accumulation and tomato gene expression, tomato plants were infected with *P. syringea* strain DC3000 isolated in 1960 in Gernsey (Buell et al. 2003) as described in (Royer et al. 2016). Leaves from control and infected plants were harvested 72h after the inoculation.

For each study, the harvested samples were frozen in liquid nitrogen, and then stored at -80°C before being crushed in a fine powder using mortar and pestle. For each sample, two aliquots (100 mg each) were prepared, for RNA and phenolic extractions, and stored at -80°C.

### qPCR analysis

Total RNA extraction was realized from 100 mg frozen tissue using the E.Z.N.A. Plant RNA kit (Omega bio-tek, Norcross, GA, USA). A on-column-DNAse treatment (Qiagen, Hilden, Germany) was included during the RNA extraction. RNA quality was checked on a 1% agarose gel. RNA samples were quantified using absorbance at 260 nm, and their purity were evaluated by the 260/280 nm absorbance ratio. A total of 200 ng of DNA-free RNA samples was reverse transcribed in a final volume of 20 µL using the RNA to cDNA kit (Life technologies). cDNAs were then diluted ten times. Tomato gene expression was analysed by qPCR on a StepOnePlus RealtimePCRSystem (ThermoFisher Scientific, Bremen, Germany) using the primers described in supplementary data 2. Each PCR reaction was realized in 20µl containing primer pairs at 0.3µM, 5µl of cDNA and 1x SYBR® Premix Ex Taq^™^ (Takara, Shiga, Japan) and followed amplification protocol consisting in a denaturing step at 95°C for 30 s, 40 cycles at 95°C for 5 s and 60°C for 30 s. A melting curve stage was included in order to confirm the specificity of each reaction. Standard curves for absolute quantification were generated from purified pCR8 vectors containing the different targeted cDNA. Concentration of the standard targeted cDNAs, expressed as the number of copies per volume unit, were calculated using their absorbance at 260 nm (for dsDNA, 1.0 A_260_ = 50 µg/ml), the average molecular weight of pCR8/targeted cDNA (1 bp = 660 Da) and the Avogadro constant (6.022*10^23^ molecules/mol). These standard solutions were serially diluted to obtain standard series ranging from 10^4^ to 5 copies per µl of standard, each step differing by 5-fold.

### Expression and purification of recombinant proteins

The four full-length sequences (*Solyc11g071470, Solyc11g071480, Solyc06g074710* and *Solyc11g066640*) were cloned from tomato cDNA using the primers listed in supplementary data 2. The PCR products were sub-cloned first in the pCR8 vector (Invitrogen, Carlsbad, California), prior to sequencing and then transferred into pET28b His-tag expression vector as a *BamH*I-*Not*I fragment. The four expression vectors were introduced into *E. coli* strain BL21-DE3 by heat shock transformation and the transformants were selected on kanamycin. Protein expression was realized by growing 50 mL of culture at 37°C until a DO_600_ comprised between 0.4 and 0.6. Then 1 mM of IPTG was added and the cultured was incubated for 22 h at 18°C. The bacteria were then harvested by a 5000 g centrifugation for 15 min at 4°C, then washed three times in 5 mL of PBS buffer (8.1 mM Na_2_HPO_4_, 1.76 mM KH_2_PO_4_, 2.7 mM KCl, 137 mM NaCl, pH 7.4) and finally resuspended in 1 mL of PBS. Bacterial lysis was realized by three 30 s runs of sonication (Bandelin Sonoplus HD 2010 MS73 probe with an intensity of 200 W/cm^2^). After a final centrifugation at 16000 g for 10 min at 4°C, the supernatant containing the protein of interest was collected and stored at -20°C. His-tag proteins were purified using a Ni-NTA purification system (Qiagen) by following the specification of the manufacturer. Purified His-tag proteins were then used directly to perform enzymatic assays.

### Enzymatic characterization

CoA esters of *p*-coumaric, caffeic, and ferulic acids were synthesized using the recombinant 4-CL enzyme from *N. tabacum*. Enzymatic assays were performed in a total volume of 100 µL containing 40µM of potential acyl donors (*p*-coumaroyl-, caffeoyl- and feruloyl-CoA) and 500µM of polyamines (putrescine, cadaverine, agmatine, spermidine, spermine, tyramine, octopamine, dopamine, noradrenaline) as acyl acceptors in 100 mM Tris-HCl buffer, pH 9, containing 5 mM EDTA. Since the amount of overexpressed enzymes made them hardly quantifiable (Supplementary data 3), the enzyme activity assays were realized, for each enzyme, with a same volume of purified enzyme (10µl) for all the substrate tested. This procedure allows comparing substrate specificity for each enzyme and the substrate preference between the four enzymes only on a qualitative basis. The reactions were initiated by the addition of purified enzymes, incubated at 35°C for 5 min, and stopped by adding 50µL of acetonitrile containing 1% HCl. The reaction mix was then centrifuged 10 min at 14000 g and filtrated on 0.2 µm filter before being analysed on UPLC-DAD-ESI-MS according to (Larbat et al. 2014). The kinetic constants of the four enzymes were determined using a range of concentration of their respective preferential acyl donor (from 5 to 60 µM for *p*-coumaroyl-CoA, caffeoyl-CoA and feruloyl-CoA) and of their preferential acyl acceptor (from 20 to 400 µM putrescine). Products were characterized according to their mass and mass fragmentation. Product accumulation was quantified by measuring the area under peak and converted with respect to caffeic, ferulic and *p*-coumaric acid standard curves. The apparent K_m_ was determined by Lineweaver-Burk extrapolation. The optimum pH was determined by measuring the caffeoyl-putrescine synthesis activity for all tomato genes, except Solyc11g066640 which was determined for the feruloyl-putrescine synthesis activity, under a pH range from 5 to 13 by using different buffers at 100mM in the reaction mix containing 40µM of acyl donor and 500µM of putrescine (Sodium phosphate from pH 5 to 8; Tris-HCl from pH 8 to 10 and Glycine from pH 10 to 13). The optimal temperature as determined on a range from 20 to 45°C.

### Phenolamide profiling by U-HPLC-MS analysis

Phenolic compounds including phenolamides were extracted from tomato plants as previously described (Royer et al. 2016). Briefly, about 100 mg FW of tissues were extracted with 1 ml 80% MeOH and blended for 30 s and centrifuged at 4000 g for 10 min. The extraction was repeated and the two supernatants were pooled in a new tube and vacuum dried. The dried residue was then dissolved in 1 mL of MeOH 70% and filtered (0.2 µm) before analyses.

A qualitative analysis of phenolics from the different tomato organs was realized on a HPLC-DAD-LTQ Orbitrap MS (ThermoFisher Scientific, San Jose, CA, USA) as described in (Larbat et al. 2014). Quantitative data were obtained by analyzing 5 µl of each extract on a U-HPLC-DAD-MS system (Shimadzu, Japan) already described in (Larbat et al. 2014). Caffeoylputrescine and feruloylputrescine concentrations were expressed as equivalent of ferulic and caffeic acids. The concentrations of other phenolamides were expressed as mass signal intensity per mg of dry weight.

## Results

### Seeking for homologous coding sequences of PHT Solyc11g071470/ Solyc11g071480 in tomato

Investigations of *Solanum lycopersicum* genomic data using BLASTp on predicted proteins revealed two homologous sequences of Solyc11g071470 and Solyc11g071480. These two sequences are Solyc06g074710 and Solyc11g066640, sharing around 65% protein sequence identity to both Soly11g071470 and Soly11g071480. Their corresponding genes are located respectively on a distant locus and a different chromosome compared to the couple *Solyc11g071470*/*Solyc11g071480* (Supplementary data 4-A). Phylogenetic analysis with representative BAHD N-hydroxycinnamoyl transferase members indicated that these 4 tomato sequences belong to the BAHD clade IV, clustering in a sub-group of enzymes involved in putrescine monoacylation, including the *N. attenuata* AT1 and the *O. sativa* PHT and PHT3 (Fig. 1). Taking into account the phylogenetic analysis of the protein sequences, the genomic organization of the genes and the genetic distance between the promoter sequences of each gene (Supplementary data 4), it can be assumed that *Solyc11g071470*/*Solyc11g071480* are duplicated orthologs of *Na*AT1, whereas *Solyc06g074710*/*Solyc11g066640* are their paralogs.

**Figure 1:**
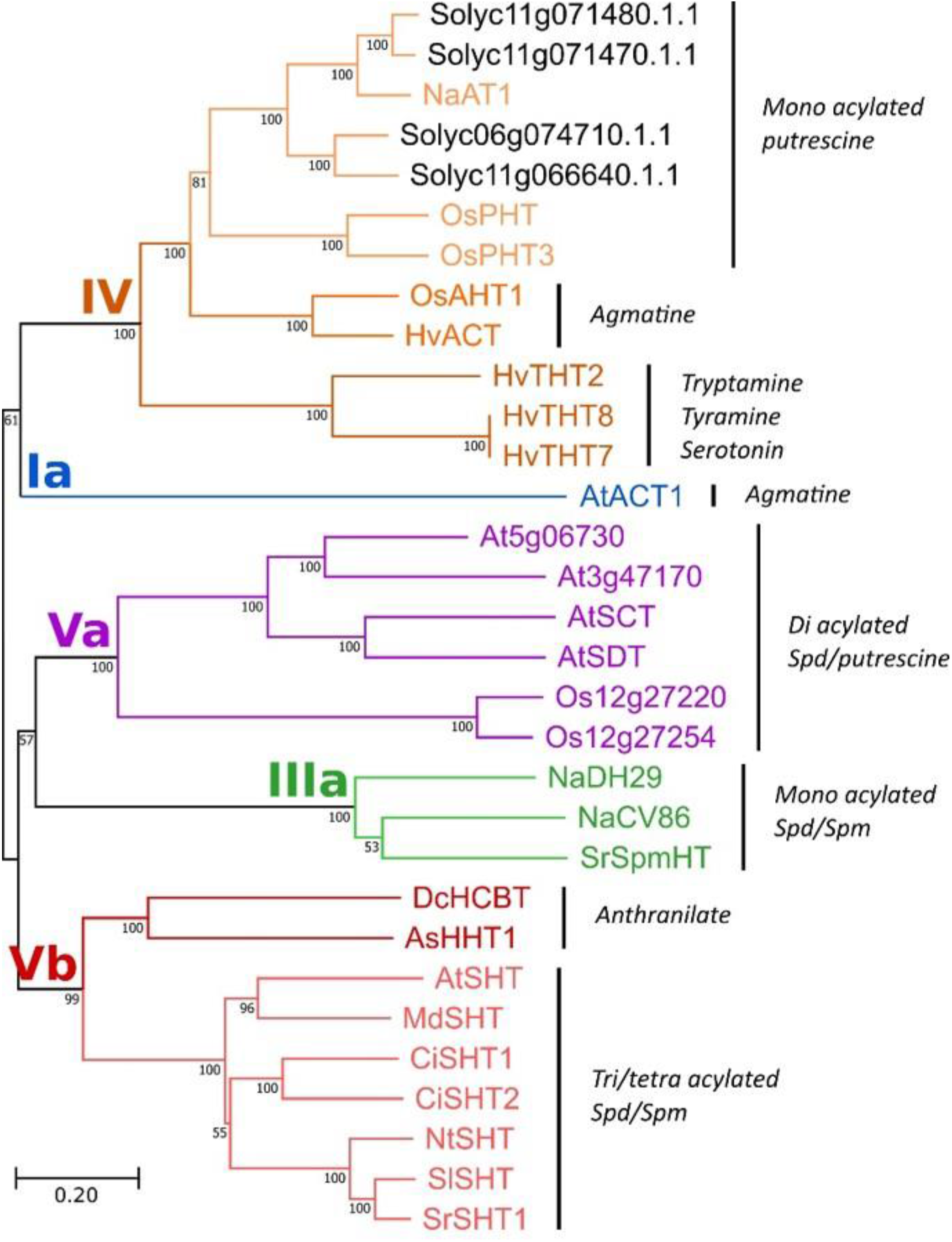
Phylogenetic tree of BAHD N-hydroxycinnamoyltransferases. The tree was constructed by neighbor-joining distance analysis on protein sequences. The putative N-hydroxycinnamoyltransferases Solyc11g066640 and Solyc06g074710 from tomato were compared to a selection of BAHD N-hydroxycinnamoyltransferases from various species: *S. lycopersicum* soly11g071470, soly11g071480 and *Sl*SHT (MN787044); *N. attenuata Na*AT1 (JN390826), *Na*DH29 (JN390824) and NaCV86 (JN390825); *O. sativa Os*PHT (BAT09226), *Os*PHT3 (BAT09225), *Os*AHT1 (BAS91492), Os12g27220 (ABA98379) and Os12g27254 (ABG22016); *H. vulgare Hv*ACT (AY228552), HvTHT7 (Hr1G019410.1), HvTHT8 (Hr1G077780.1) and HvTHT2 (Hr1G077790.1); *A. sativa As*HHT1 (AB076980); *A. thaliana At*SHT (NM127464), *At*SDT (NM127915), *At*SCT (NM128072), *At*ACT1 (BT011800), At5g06730 and At3g47170; *S. richardii Sr*SHT (KP165411) and *Sr*SpmHT (KR150683); *D. caryophyllus Dc*HCBT (Z84383); *M. domestica Md*SHT (ALF00095); *C. intybus Ci*SHT1 and 2 (MG457243, MG457244); *N. tabacum Nt*SHT (MN787045).

### Enzymatic characterization

The enzymatic activity of Solyc06g074710 and Solyc11g066640 was assessed *in vitro* on purified recombinant protein expressed in *E*.*coli*. As expected, each enzyme was proved functional and exhibited a putrescine hydroxycinnamoyl-CoA transferase activity. For this reason, and for the ease of the reading, we attributed the name *Solanum lycopersicum* putrescine hydroxycinnamoyl transferase (*Sl*PHT) to each enzyme with the following nomenclature *Sl*PHT1 (Solyc11g071470), *Sl*PHT2 (Solyc11g0711480), *Sl*PHT3 (Solyc06g074710) and *Sl*PHT4 (Solyc11g066640). In our *in vitro* conditions, *Sl*PHT3 transferred efficiently caffeoyl-CoA, feruloyl-CoA and *p*-coumaroyl-CoA to both putrescine and agmatine (Fig. 2 A-B; Supplementary data 5). In addition, this enzyme also transferred caffeoyl-CoA and *p*-coumaroyl-CoA to spermidine with a lower efficiency and led, at least, to isomers of monosubstituted spermidine (Fig. 2 B; Supplementary data 5). The second enzyme, *Sl*PHT4, exhibited a narrow specificity to feruloyl-CoA which could be transferred on putrescine, agmatine and, to a lower extent, to spermidine (Fig. 2 A-B; Supplementary data 5). These results make evidence that both enzymes had different enzymatic specificities than *Sl*PHT1 and *Sl*PHT2 that exhibited a higher affinity for caffeoyl-CoA (Fig. 2 A-B-C; Supplementary data 5). The four *Sl*PHTs displayed optimal pH at 9 (*Sl*PHT1-2) and 10 (*Sl*PHT3-4). Interestingly, *Sl*PHT1-2 exhibited a significant activity on a broader range of pH from 5 to 11 (Supplementary data 6). They all had a maximum activity between 30 and 40°C, precisely 40°C for *Sl*PHT1-2, 30°C for *Sl*PHT3 and 35° for *Sl*PHT4 (Supplementary data 7). In addition to the polyamines cited above, these enzymes were also assayed with spermine, tyramine, octopamine, noradrenaline, dopamine and lysine as acyl acceptors, based on the occurrence of related phenolamides in tomato (Von Roepenack-Lahaye et al. 2003; Zacarés et al. 2007; Larbat et al. 2014), however no activity was detected.

**Figure 2:**
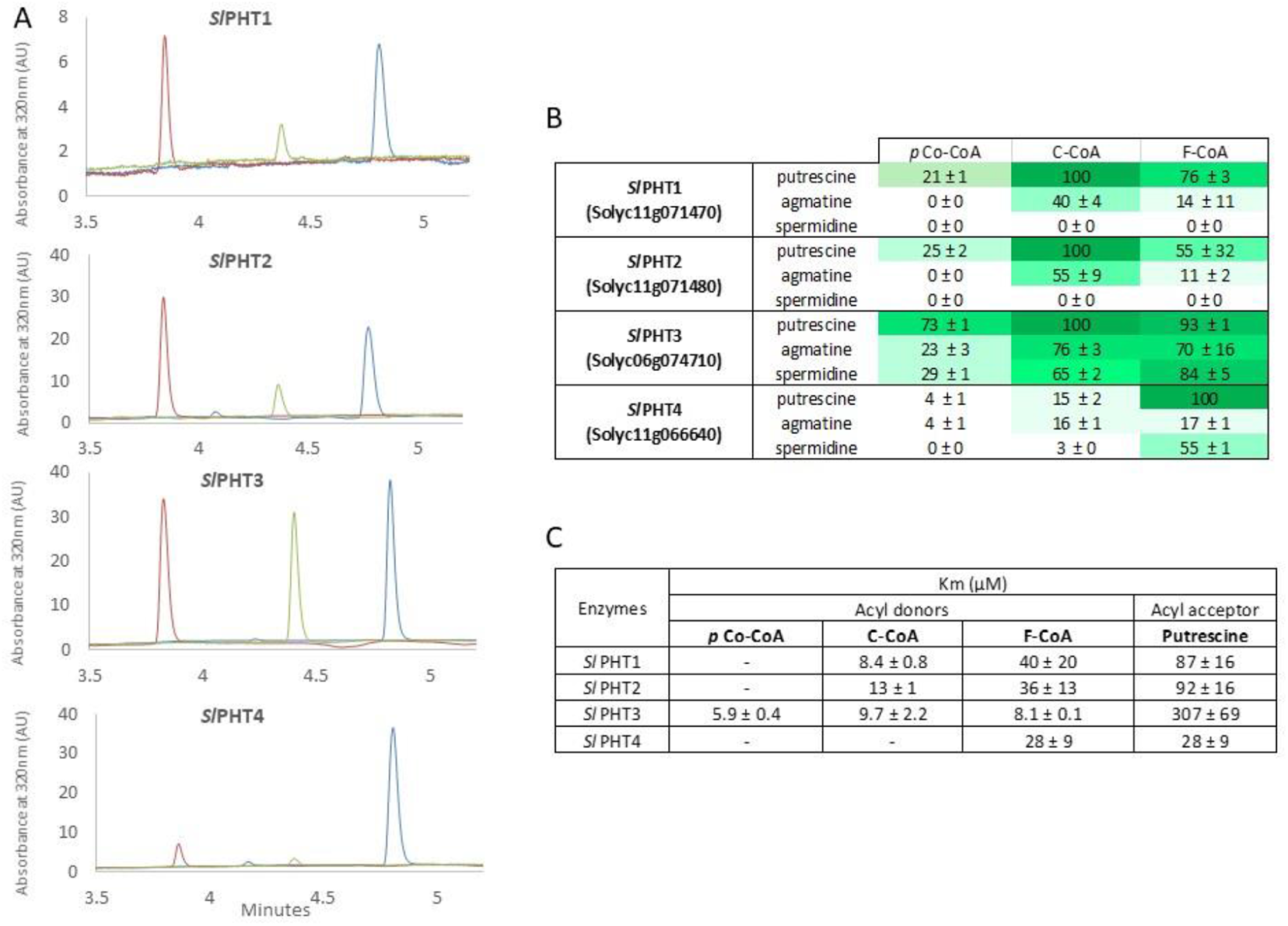
Functional characterization of putrescine hydroxycinnamoyltransferases from tomato. A: Chromatograms showing the amount of *p*-coumaroyl putrescine (green), caffeoyl putrescine (red) and feruloylputrescine (blue) produced from independant reactions with the same reaction conditions and enzyme amount for the 4 tomato hydroxycinnamoyltransferases. B: Substrate specificity of the four enzymes. *p*-coumaroyl-CoA (pCo-CoA), caffeoyl-CoA (C-CoA) and feruloyl-CoA (Fer-CoA) were incubated with the respective acyl acceptors (putrescine, agmatine and spermidine). The amount of product formed per time unit was determined by measuring the area under peak at 320 nm and expressing as *p*-coumaric acid, caffeic acid and ferulic acid equivalents. Then, substrate preference was expressed relative to the best reaction. The color scale indicates the favorable reactions in green. C: Catalytic constants determined for putrescine and the three main acyl donors.

### Phenolamide distribution in tomato

Caffeoyl putrescine and feruloyl putrescine were the two main phenolamides accumulating in vegetative organs of plantlets and 28-day old tomato plants (Fig. 3 A). Both metabolites were also found in flowers although these organs contained a much higher diversity of phenolamides, the major ones being di- and tri-coumaroyl spermidines (Supplementary data 8). Caffeoyl putrescine was mainly detected in the higher part of stems with a concentration reaching 5.5 ± 1.3 µg.g FW^-1^. In the other plant parts, the concentration of these molecules was below 1 µg.g FW^-1^. The distribution of feruloyl putrescine was different than caffeoyl derivatives. It mostly accumulated in roots from plantlet (0.4 ± 0.04 µg.g FW^-1^, in roots of 28-day old plants (1.2 ± 0.1 µg.g FW^-1^) and in flowers (from 1.3 ± 0.2 to 2.3 ± 0.7 µg.g FW^-1^, Fig 3 A).

**Figure 3:**
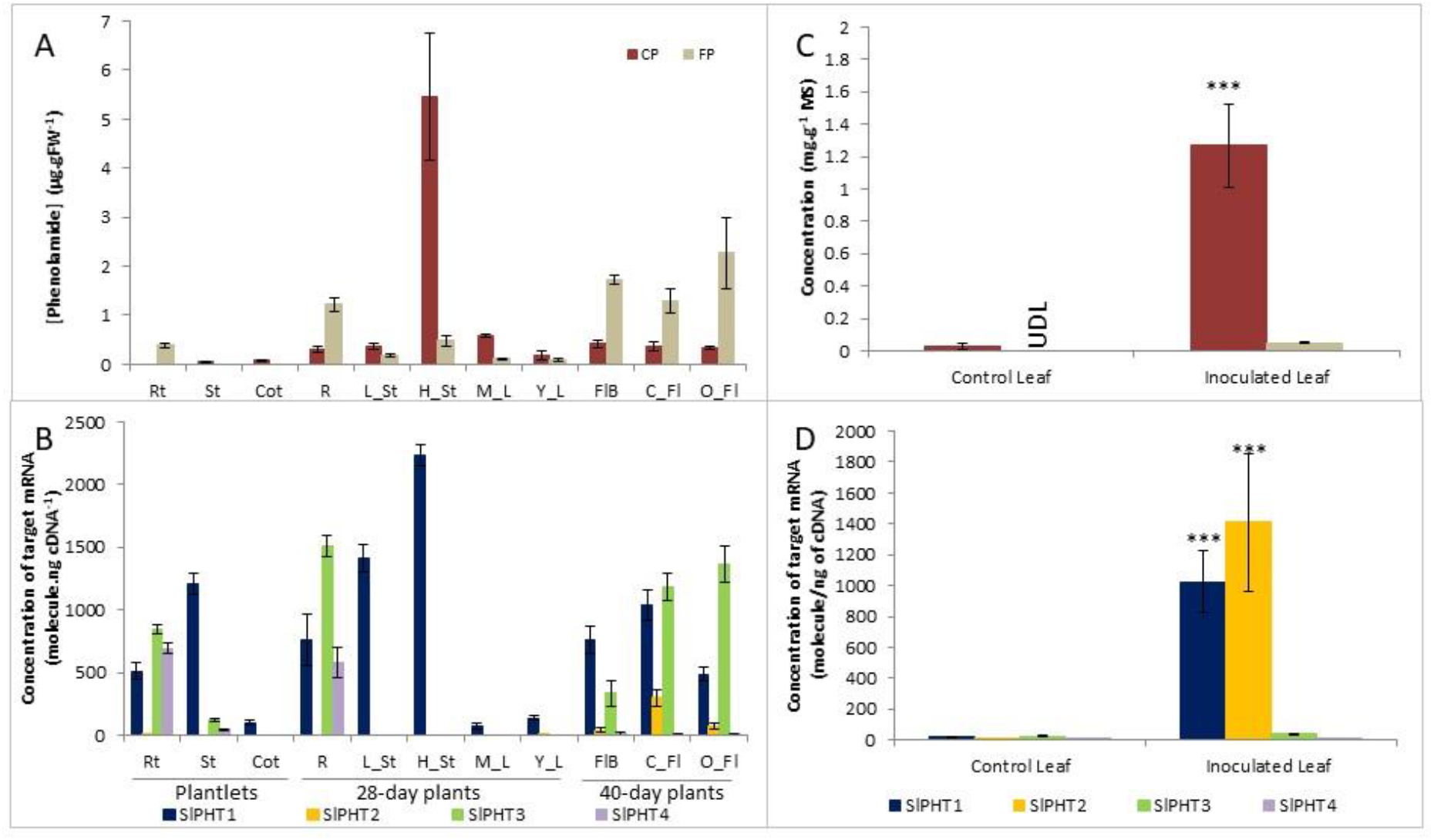
Regio-specific and inducible accumulation of putrescine-associated phenolamides and the 4 *Sl*PHT transcript abundance in tomato. The concentrations of caffeoylputrescine (CP) and feruloylputrescine (FP) were determined in different tomato organs (A) and in tomato leaves inoculated with *Pseudomonas syringae* for 72h (B). The absolute expression levels of the 4 genes coding for *Sl*PHT were measured in the same tomato organs (C) and in tomato leaves inoculated with *Pseudomonas syringae* for 24h (D). Error bars represent the standard errors calculated from four biological replicates. Rt: Rootlet; St: Plantlet Stem; Cot: Cotyledon; R: Root; M_L: Mature Leaves; Y_L: Young Leaves; FlB: Flower Buds; C_Fl: Closed Flower; O_Fl: Open Flower. L_St: Lower Stem; H_Stem: High Stem; UDL: Under Detection Limit. P<0.001 *** T. test.

As already reported (Royer et al. 2016), we confirmed that the foliar accumulation of caffeoyl putrescine was dramatically increased upon infection with *Pseudomonas syringae* (42-fold increase in the infected leaf compared to the control; Fig. 3 B). The infection with *P. syringae* induced also the foliar accumulation of feruloyl putrescine that was not detected in the control leaf (Fig. 3 B).

### Gene expression in planta

Using the same set of tissues as for metabolic analysis, we measured the gene expression of the four *Sl*PHT using quantitative RT-PCR (Fig. 3 C). *Sl*PHT1 gene expression was detected in all plant parts. In the vegetative organs, the expression level was the highest in stems, then in roots and, to a lesser extent, in cotyledons and leaves. Interestingly, *Sl*PHT1 was the only gene which expression was detected in the aboveground tissues of the 28-day old plants. *Sl*PHT3 and 4 displayed a similar expression pattern and are restricted to roots and rootlets. Finally, *Sl*PHT2 expression was not detected in any vegetative tissues. In flowers, we could show the expression of all four *Sl*PHTs. *Sl*PHT1 was expressed with the same order of magnitude in flower buds and opened flowers, whereas the expression level of *Sl*PHT3 was increasing according to the development stage of the flowers. Finally, although detected, the expression level of *Sl*PHT2 and 4 in flowers was low.

The inoculation of tomato leaves with *P. syringae* led to a strong modification of the expression pattern of the PHT genes. Indeed, while only *Sl*PHT1 was detected in control leaves, the expression of all *Sl*PHT genes but *Sl*PHT3 was detected in inoculated leaf. Among them, *Sl*PHT2 and *Sl*PHT1 were markedly induced (51- and 809-fold increase respectively, Fig. 3 C)

## Discussion

Despite their wide distribution among plants, their high structural diversity and their involvement in several biological processes, the understanding on the phenolamide metabolism and regulation is far from being complete. Our work provided insights on putrescine derivatives accumulation in tomato plants by the identification and characterization of two new N-hydroxycinnamoyltransferases. These two enzymes, in addition to the other two enzymes that we have previously reported (Roumani et al. 2022), belong to a phylogenetic cluster of putrescine hydroxycinnamoyltransferases. The cluster contains *Na*AT1 putrescine transferase isolated from *N*.*attenuata* (Kaur et al. 2010). The two new identified enzymes are close to the rice transferases that uses putrescine and agmatine as acyl acceptors to synthesize *p*-coumaroyl and feruloyl derivates (Tanabe et al. 2016).

The functional characterization together with the expression pattern *in planta* of the four *Sl*PHT indicated that the synthesis of putrescine containing phenolamides in tomato is controlled by a family of genes sharing a complementarity in function and spatial expression. This conclusion is in accordance to the observations made on rice (*O. sativa*) where the root and flower accumulation of agmatine and putrescine phenolamides were correlated to the spatial expression of three functionally characterized PHT (Tanabe et al. 2016). In our study, the combined analysis of substrate specificity and spatial expression allow us to assume that the above-ground accumulation of caffeoylputrescine is principally controlled by *Sl*PHT1. In roots, the high accumulation of feruloylputrescine correlates with the expression of *Sl*PHT4 which exhibited a narrow specificity to feruloyl-CoA, and also the expression of *Sl*PHT3 which had a wider range of substrates. In flowers, the accumulation of feruloylputrescine is not correlated to the *Sl*PHT4 expression, but could be related to the expression of *Sl*PHT2 and *Sl*PHT3. Additional mechanisms may however contribute to better explain the specific ratio found between caffeoyl and feruloyl derivatives in the different plant parts, of which substrate availability appears as a major one. Moreover, the accumulation of other phenolamides in flowers could be partially explained by the specificity of *Sl*PHTs. Indeed, the ability of *Sl*PHT3 to catalyse, *in vitro*, the synthesis of monoacylated spermidine, suggests that it could contribute to the accumulation of *p*-coumaroylspermidine and feruloylspermidine. Since the specific accumulation of trisubstituted spermidine derivatives in flowers is controlled by another subclass of transferase named spermidine hydroxycinnamoyl transferase (SHT, Perrin et al., 2021), the question remains regarding the accumulation of disubstituted spermidine derivatives like di-*p*-coumaroylspermidine, diferuloylspermidine or *p*-coumaroyl,feruloylspermidine (Supplementary data 8). The accumulation of such compound would require another transferase activity that has not been discovered yet. Interestingly, the expression patterns that we reported on the four *Sl*PHT were confirmed and complemented by public gene expression databases such as Tomato Expressed Database (TED, http://ted.bti.cornell.edu/cgi-bin/TFGD/digital/home.cgi) and Tomato Expression Atlas (TEA, https://tea.solgenomics.net/). With respect to these databases, *Sl*PHT3 appears as the main expressed PHT in the fruit tissues.

In accordance to their role in plant defense, phenolamides and among them, caffeoylputrescine and feruloylputrescine were significantly accumulated tomato leaves infested by *Pseudomonas syringae*. This accumulation appears correlated to the highly inducible expressions of *Sl*PHT1 and moreover *Sl*PHT2 which was barely detected in non-infested tissues. The high inducibility of these two genes was also observed in tomato leaves infested with larvae of the lepidoptera *Tuta absoluta* (Roumani et al. 2022). In addition, the inducibility of only these two *Sl*PHT was demonstrated recently in tomato leaves treated with jasmonic acid. Other phytohormones (Ethylene, salicylic acid) or the two elicitors flg15 and BP178 were however ineffective (Montesinos et al., 2021 supplementary table 2A).

In conclusion, the present study allowed the identification and the biochemical characterization of a small family of hydroxycinnamoyltransferases implicated in the biosynthesis of putrescine-containing phenolamides. The substrate specificity and the expression pattern of these four enzymes supported the spatial distribution of caffeoyl and feruloylputrescine distribution *in planta* in both healthy and infected plant.

## Author contribution statement

RL conceived the project and designed experiments. MR and RL performed the experiments. MR and RL analyzed the data. MR, SB, AH and RL prepared Figures and Tables and wrote the manuscript. All authors have approved the final version.

## Acknowledgements

The authors warmly thank Aude Fauvet, Clément Charles, Claude Gallois, Leonor Duriot, Hadjara Saindou and Krissie Loisy for their precious help regarding different aspects of the project, including plant culture and sampling, metabolite and RNA extraction and biochemical characterization of enzymes. Plants were grown on the Plant Experimental Platform in Lorraine (Université de Lorraine) and metabolomic analyses were conducted on the Metabolomic and Structural Analytic Platform (Université de Lorraine).

## Data availability

The sequences of Solyc11g071470, Solyc11g071480, Solyc06g074710 and Solyc11g066640 are available at the ncbi website with the following ID 2573112 ; 2573120 ; 2615170 and 2615163

