## Supplemental data 1 and 2 for "Functional characterization of a small gene family coding for putrescine hydroxycinnamoyltransferases in tomato"

**Supplementary Data 1: Medium composition for the hydroponic tomato culture**

A- Final concentrations of the pure salts composing the medium

| | $Ca(NO_3)_2$ | $KH_2PO_4$ | $K_2SO_4$ | $MgSO_4$ | $CaSO_4$ | EDTA-Fe | Micro-elements (*) |
|---|---|---|---|---|---|---|---|
| Concentration (mM) | 3.5 | 1.0 | 1.0 | 1.5 | 3.0 | 0.043 | 0.3 $mL.L^{-1}$ |

B- Elemental composition of the micro-element solution

| Micro-elements composition | Mo | Mn | Zn | Cu | B | Fe |
|---|---|---|---|---|---|---|
| Concentration (mM) | 0.94 | 38.8 | 10.8 | 1.6 | 68.7 | 35 |

### Supplementary Data 2 : List of primers

| Genes | Forward primer | Reverse primer | Application |
|---|---|---|---|
| Solyc11g071470 (*Sl*PHT1) | CTGTAGACGCCTCGTTGGAC | AGGAGAGCATAGAGGGAGAAGG | qPCR |
| Solyc11g071480 (*Sl*PHT2) | TATCGAGAGTGGGCAGGAAG | TCCAATGAGGTGTCCACAGA | qPCR |
| Solyc06g074710 (*Sl*PHT3) | AGGCCCTAGCGATTTACAGG | CCTCAACGAATCGAACACCT | qPCR |
| Solyc11g066640 (*Sl*PHT4) | GCGGGCTAAGTGGATTTGA | GTGTGGCAGAGGTTCTCGTAA | qPCR |
| Solyc11g071470 (*Sl*PHT1) | ACAACGGATCCATGAATGTGAAAATTGAGAGTTCAAAAATC | ACAAGCGGCCGCTCACTTTGCTTTCAAATCTA | cloning |
| Solyc11g071480 (*Sl*PHT2) | ACAACGGATCCATGAATGTGAAAATTGATAGTTCAAAAATC | ACAAGCGGCCGCTCACTTTGCTTTCAAATCTA | cloning |
| Solyc06g074710 (*Sl*PHT3) | ACAACGGATCCATGAAGGTCAAAATAGAAAGTTCAAAAATC | ACAAGCGGCCGCTCAAGCAAGATCTAAAGAATAG | cloning |
| Solyc11g066640 (*Sl*PHT4) | ACAACGGATCCATGAAGATTAAAATAGAAAGTTCAAGAATT | ACAAGCGGCCGCTTAATCTTCAAGCAAGTCCAAGGAATAA | cloning |
