## Supplementary figures and images for "Functional characterization of a small gene family coding for putrescine hydroxycinnamoyltransferases in tomato"

### Supplemental data 3

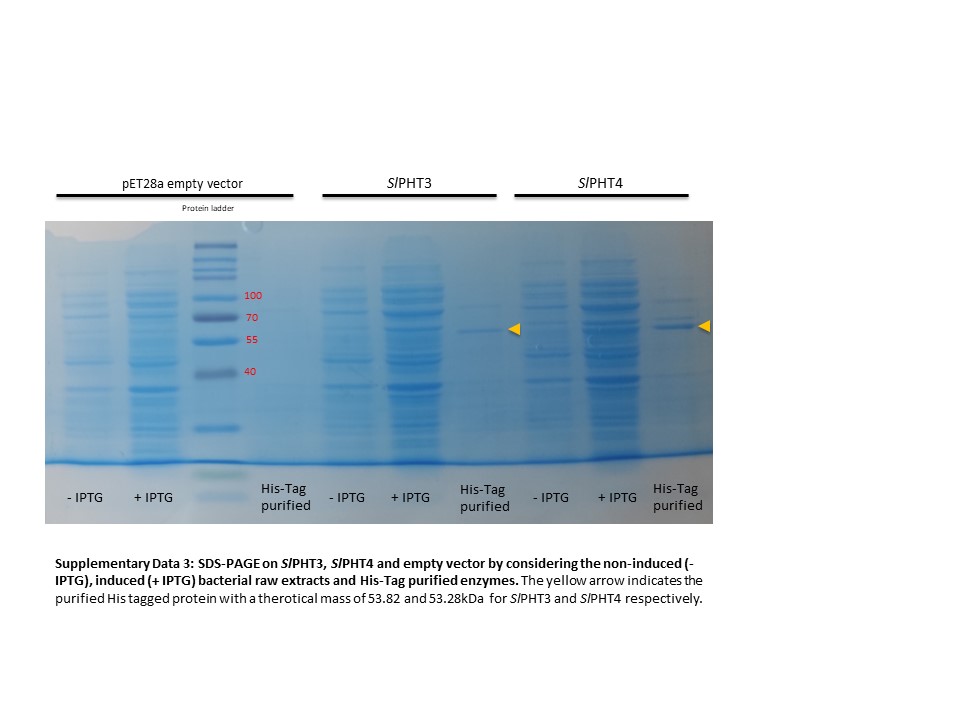

### Supplemental data 4

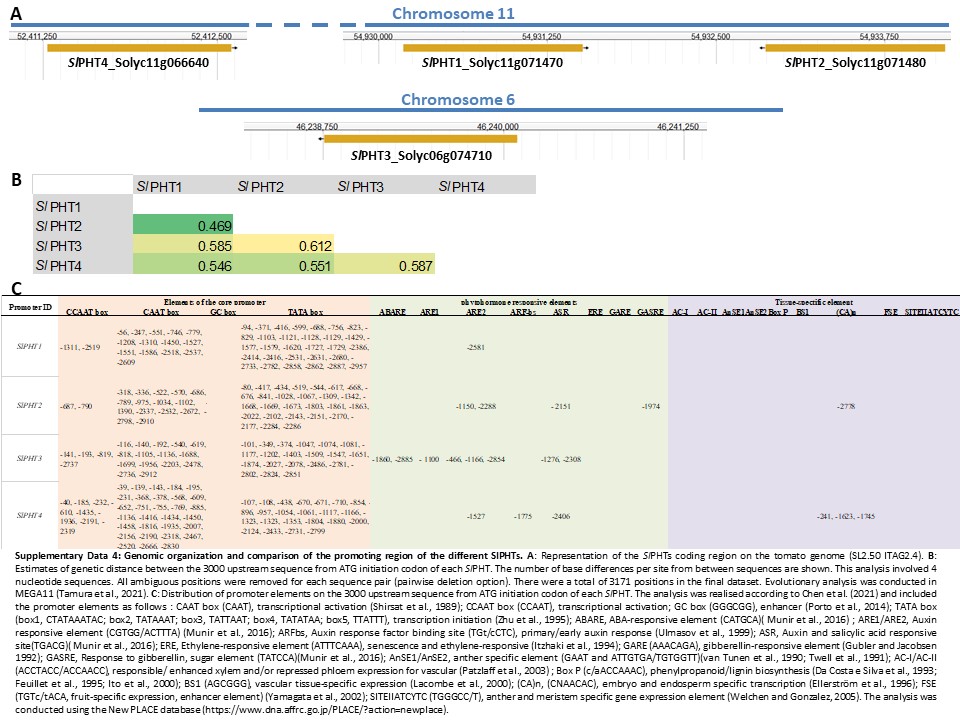

### Supplemental data 5

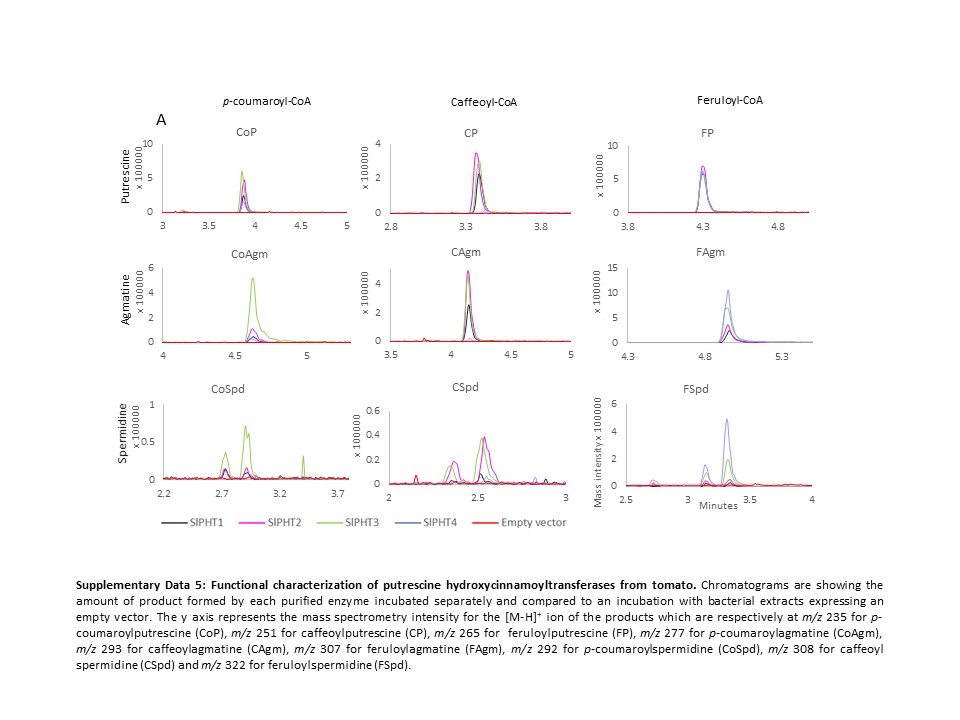

### Supplemental data 6

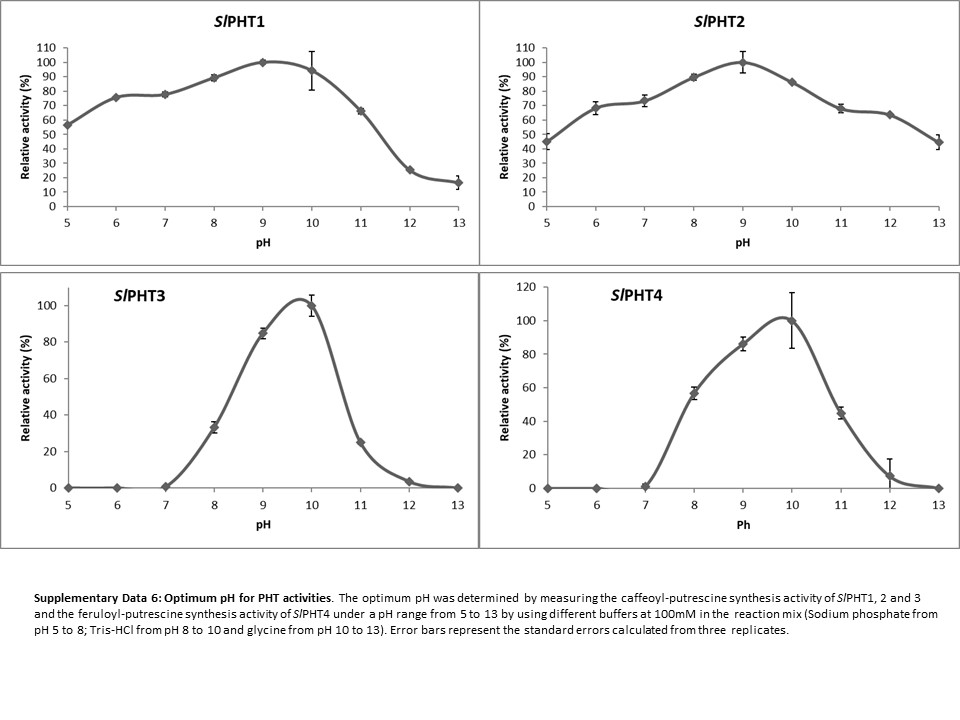

### Supplemental data 7

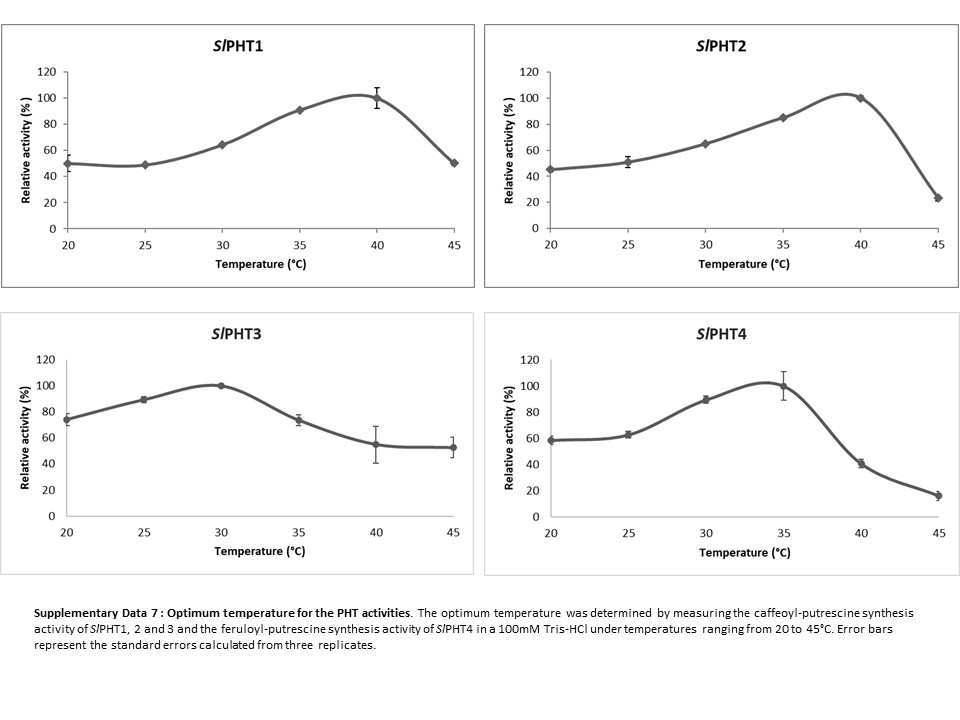

### Supplemental data 8

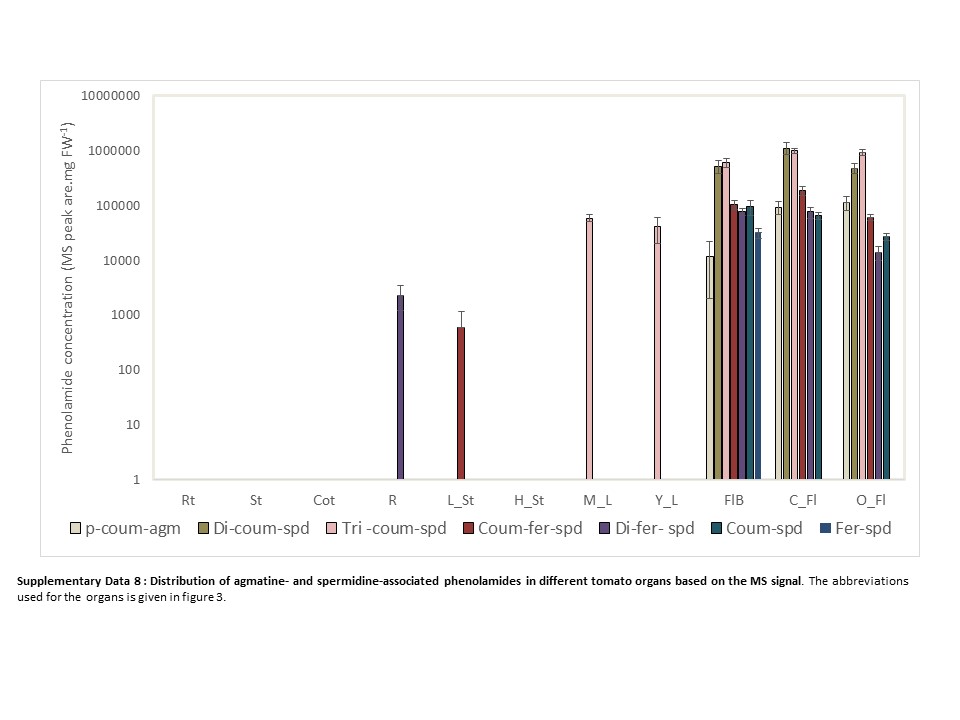
